# RawTools: Rapid and Dynamic Interrogation of Orbitrap Data Files for Mass Spectrometer System Management

**DOI:** 10.1101/418400

**Authors:** Kevin A. Kovalchik, Shane Colborne, Sandra Spencer, P.H. Sorensen, David D. Y. Chen, Gregg B. Morin, Christopher S. Hughes

**Affiliations:** Department of Chemistry, University of British Columbia, Vancouver, British Columbia, Canada; Canada’s Michael Smith Genome Sciences Centre, British Columbia Cancer Agency, Vancouver, British Columbia, Canada; Department of Molecular Oncology, British Columbia Cancer Research Centre, Vancouver, British Columbia, Canada; Department of Medical Genetics, University of British Columbia, Vancouver, British Columbia, Canada

**Keywords:** RawQuant, Orbitrap, Mass Spectrometry, Instrument Diagnosis, Isobaric Tag Quantification, Label-free Quantification, Method Optimization

## Abstract

Optimizing the quality of proteomics data collected from a mass spectrometer (MS) requires careful selection of acquisition parameters and proper assessment of instrument performance. Software tools capable of extracting a broad set of information from raw files, including meta, scan, quantification, and identification data are needed to provide guidance for MS system management. In this work, direct extraction and utilization of these data from Thermo Orbitrap raw files is demonstrated using the RawTools software. RawTools is a standalone tool for extracting meta and scan data directly from raw MS files generated on Thermo Orbitrap instruments. RawTools generates summarized and detailed plain text outputs after parsing individual raw files, including scan rates and durations, duty cycle characteristics, precursor and reporter ion quantification, and chromatography performance. RawTools also contains a diagnostic module that includes an optional ‘preview’ database search for facilitating informed decision-making related to optimization of MS performance based on a variety of metrics. RawTools has been developed in C# and utilizes the Thermo RawFileReader library, and thus can process raw MS files with high speed and high efficiency on all major operating systems (Windows, MacOS, Linux). To demonstrate the utility of RawTools, extraction of meta and scan data from both individual and large collections of raw MS files was carried out to identify problematic characteristics of instrument performance. Taken together, the combined rich feature-set of RawTools with the capability for interrogation of MS and experiment performance makes this software a valuable tool for proteomics researchers.

## Introduction

The rapid development of mass spectrometry (MS) as a tool to study and characterize the proteome has led to the creation of vast amounts of data of various qualities and levels of information content. Initially, making sense of these data was primarily addressed using software focused on the robust and accurate identification of tandem mass spectrometry (MS2) fragmentation spectra of peptides^1^, and subsequent quantification of their signal intensities to determine abundance^2^. Continued development efforts have resulted in a collection of search and quantification tools capable of performing a wide variety of analyses on data originating from diverse MS platforms. Recently, more focus has been on the development of tools aimed at monitoring the performance of MS hardware via interrogation of data files generated from analysis of standard samples^3,4^. These quality control (QC) software tools provide valuable insight into MS metrics that includes peptide identification rates, ion signal, and chromatography performance, which can all be used to temporally monitor instrument performance. However, there remains a need for software tools aimed at diagnostic management of MS systems towards improving the quality and maximizing the information content of generated data.

Parsing of meta, scan, and quantification data from a raw MS file can allow informed decision making related to the optimization of instrument and experiment performance and design. Recently, there has been active development in the parsing of these acquisition metrics from raw MS data^5–7^. Acquisition data metrics can include information such as scan rate, duty cycle time, ion injection times, and number of triggered dependent events per cycle. These metrics are useful for optimization of MS method parameters and in the provision of greater insight during the tracking of instrument performance. Unfortunately, obtaining these data from raw files using non-commercial and non-vendor provided software is made challenging due to the proprietary nature of the raw MS data file format. To facilitate these types of analyses, we previously developed the RawQuant tool. RawQuant enabled parsing of meta and scan data, in addition to quantification of isobaric tag reporter ions directly from Orbitrap raw MS files^8^. Using RawQuant, the ability to efficiently identify key areas where MS performance was suboptimal was demonstrated, along with the limited utility of a variety of isobaric tag quality filtering approaches^8^. Importantly, RawQuant was provided as an open-source and freely available tool that generated easily parsed plain text output from raw Orbitrap MS file contents. Although efficient at processing raw Orbitrap files for meta, scan, and quantification data, RawQuant was built around the Thermo MSFileReader library that rendered it incompatible with non-Windows operating systems. In addition, the development of RawQuant in Python necessitated numerous installation steps and limited processing performance.

In this work, we present the development of the RawTools software package as a substantial improvement over RawQuant with improved speed, increased functionality, and expanded operating system compatibility. RawTools has been built from the ground up in C# to facilitate easier implementation on user machines, as well as to provide significant processing speed gains. RawTools maintained much of the functionality of its predecessor, such as parsing of meta and scan data along with isobaric tag quantification, but has also incorporated new features that include label-free parent ion quantification and direct assessment of chromatography performance. In addition, RawTools contains a newly developed ‘QC’ functionality for diagnostic analysis using a large number of summarized metrics to facilitate informed decision-making towards achieving and maintaining optimum MS performance. As part of the QC feature, RawTools also directly communicates with the open-source and freely available search tool IdentiPy to perform a rapid ‘preview’ search of the processed data. This preview search enables tracking of metrics that include proteolysis performance, isobaric labeling efficiency, and MS2 identification rates. Importantly, diagnostic QC analysis with RawTools allows rapid instrument parameter optimization and experiment quality monitoring in near real-time due to the speed and efficiency of the software tool. Moreover, diagnostic results can be easily dissected by any user using the newly developed R Shiny application web interface for RawTools. Lastly, RawTools has been built to utilize the Thermo RawFileReader package, enabling universal direct processing of Orbitrap raw files on Windows, MacOS, and Linux platforms. Like RawQuant, RawTools is open source, freely available, and includes detailed step-by-step and image-rich user documentation to ensure maximum usability by users of all skill levels. Altogether, these features make RawTools a valuable software tool for the rapid and dynamic processing of raw data acquired on Thermo Orbitrap MS instruments towards performance optimization and monitoring to ensure the consistent acquisition of high-quality and information-rich data.

## Experimental Section

### RawTools software, documentation, and availability

RawTools was developed in the C# programming language and is covered by the Apache 2.0 license. C# is a .NET language, which makes it natively compatible with the RawFileReader library. The RawFileReader library is developed and distributed by Thermo Scientific under its own license, and is separate from RawTools. RawFileReader is built on the .NET framework, which is compatible with all three major operating systems (e.g. Windows, MacOS, Linux). RawTools utilizes RawFileReader in order to access the meta and scan data present in the raw MS files. RawTools was developed based on the use of the .NET framework version 4.6.2 or greater, facilitating support across a wide range of Windows operating systems. For use on Linux and MacOS machines, RawTools was extensively tested using Mono (version 5.12.0.233), which clones the .NET framework for use on Unix systems.

RawTools is open source and freely available. The code and compiled versions along with step-by-step walkthroughs and image-rich documentation are available on GitHub: https://github.com/kevinkovalchik/RawTools. In lieu of providing supplementary tables with this work describing the RawTools output, detailed parameter-by-parameter documents are available on the GitHub page. As RawTools will be in active development for the foreseeable future, the software and supporting documentation are in constant flux. Therefore, the most up-to-date versions can always be found in the publicly accessible GitHub resource. Lastly, an R Shiny application developed to enable direct visualization and interpretation of RawTools QC results is freely available on the web at: https://rawtoolsqcdv.bcgsc.ca. The RawTools R Shiny application is also freely available on the GitHub page if use on a local machine is preferred: https://github.com/kevinkovalchik/RawTools/tree/master/documentation/manuscript/RawTools_RShiny_Application. This web application requires the comma-separated output from a RawTools diagnostic analysis (‘QC’ function in RawTools (Figure 1)) analysis. Example diagnostic QC data that can be used with the R Shiny application is available for free download on the RawTools GitHub page (https://github.com/kevinkovalchik/RawTools/tree/master/documentation/QC_example-data).

**Figure 1.**
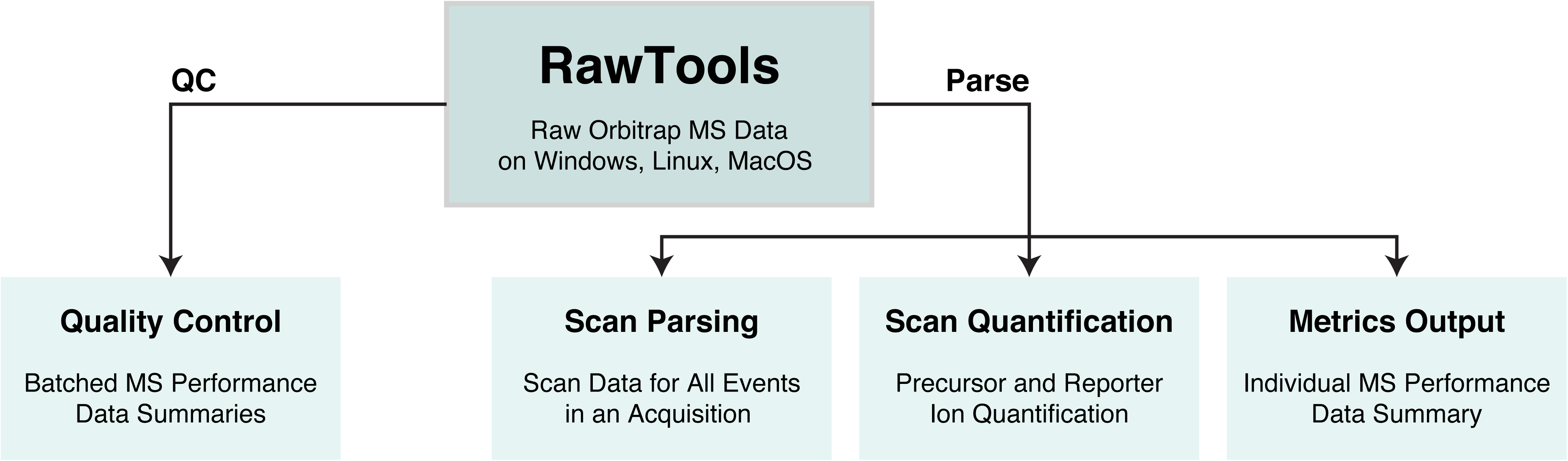
RawTools includes a wide range of built-in utilities and potential functionality for raw MS data processing. Schematic depicting the different functional modules of RawTools. The software is divided into ‘parse’ and ‘qc’ processing pipelines that generate overlapping, but individual data sets as indicated. All functionalities displayed work directly with raw MS files derived from Thermo Orbitrap instruments on Windows, Linux, and MacOS computational hardware.

### Cell culture and harvest

HeLa cells were grown and harvested by the National Cell Culture Center (Biovest International). A total of 1×10^9^ cells were grown and provided as aliquoted pellets at a concentration of 5 × 10^7^ cells per tube. Cells were stored at -80°C until use.

### Guanidine-based protein isolation, reduction, alkylation, and digestion

Cell pellets (5 × 10^7^ cells each) were thawed on ice with periodic vortexing. To each pellet, 900μL of lysis buffer (50mM HEPES pH 8, 4M guanidine hydrochloride (Sigma, CAT#G3272), 50mM NaCl (Sigma, CAT#S7653), 10mM tris(2-carboxyethyl)phosphine hydrochloride (TCEP) (Sigma, CAT#C4706), 40mM chloroacetamide (CAA) (Sigma, CAT#C0267), 1X cOmplete protease inhibitor – EDTA free (Sigma, CAT#11836170001)) was added, and the pellets mixed using an 18G syringe. Lysis mixtures were transferred to 2mL FastPrep-compatible tubes containing Lysing Matrix Y (MP Biomedicals, CAT#116960050). Lysis mixtures were vortexed on the FastPrep-24 5G instrument (MP Biomedicals) (6 M/s, 40 seconds, 2 cycles, 120 second rest between cycles). Lysates were centrifuged at 20,000g for 5 minutes and the supernatant recovered. Resultant lysates were heated at 65°C for 15 minutes, and chilled to RT for a further 15 minutes. Protein concentrations were approximated using A280 readings from a NanoDrop instrument (Thermo Scientific).

For in-solution digestion, 200μg of HeLa protein sample was diluted 1:10 with 50mM HEPES pH 8, and trypsin/rLysC mix (Promega, CAT#V5071) (1:25 enzyme to protein concentration) was added prior to incubation for 14 hours at 37°C in a ThermoMixer with mixing at 1000rpm (final digestion volume per sample ≈400μL). Following digestion, mixtures were acidified by addition of trifluoroacetic acid (TFA) (Sigma, CAT#302031) to a 1% final concentration. Tubes were centrifuged at 20,000g for 1 minute to pellet any precipitate and the supernatant was recovered for further processing.

### Peptide clean-up

Peptides were desalted and concentrated using TopTips (Glygen). For TopTip clean-up, 1mL TopTips (CAT#TT3C18) were rinsed twice with 0.6mL of acetonitrile (Sigma, HPLC-grade, CAT#34998) with 0.1% TFA. Cartridges were then rinsed twice with 0.6mL of water (Sigma, HPLC-grade, CAT#34877) with 0.1% TFA prior to sample loading. Loaded samples were rinsed three times with 0.1% formic acid (0.6mL per rinse) and eluted with 1.2mL of 80% acetonitrile containing 0.1% formic acid. Eluted peptides were concentrated in a SpeedVac centrifuge (Thermo Scientific) and subsequently reconstituted in 1% formic acid (Thermo Scientific, LC-MS grade, CAT#85178) with 1% dimethylsulfoxide (Sigma, CAT#D4540) in water.

### Mass spectrometry data acquisition

Analysis of peptides was carried out on an Orbitrap Velos MS platform (Thermo Scientific). Samples were introduced using an Easy-nLC 1000 system (Thermo Scientific). The Easy-nLC 1000 system was plumbed with a single-column setup using a liquid-junction for spray voltage application. The factory 20µm ID × 50cm S-valve column-out line was replaced with a 50µm ID x 20cm line to reduce backpressure during operation at high flow rates. The column used for analytical separations was packed in-house in a 200µm ID capillary that was prepared with a fritted nanospray tip (formamide and Kasil 1640 in a 1:3 ratio) using a laser puller instrument (Sutter Instruments). The 200µm ID analytical column was packed to a length of 20cm with 1.9µm Reprosil-Pur C18 beads (Dr. Maisch) in an acetone slurry. The analytical column was connected to the Orbitrap Velos MS using a modified version of the UWPR Nanospray source (http://proteomicsresource.washington.edu/protocols05/nsisource.php) combined with column heating to 50°C using a 15cm AgileSLEEVE column oven (Analytical Sales & Service). Prior to each sample injection, the analytical column was equilibrated at 400bar for a total volume of 4μL. After injection, sample loading was carried out for a total volume of 8μL at a pressure of 400 bar. After loading, elution of peptides was performed with a gradient of mobile phase A (water and 0.1% formic acid) from 3 – 7% B (acetonitrile and 0.1% formic acid) over 2 minutes, to 30% B over 24 minutes, to 80% B over 0.5 minutes, hold at 80% B for 2 minutes, to 3% B in 0.5 minutes, and holding at 3% for 1 minute, at a flow rate of 1.5μL/min.

Data acquisition on the Orbitrap Velos (control software version 2.6.0.1065 SP3) was carried out using a data-dependent method with MS2 in the ion trap. The Orbitrap Velos was operated with a positive ion spray voltage of 2400 and a transfer tube temperature of 325°C. Survey scans (MS1) were acquired in the Orbitrap at a resolution of 30K, across a mass range of 400 – 1200 m/z, with an S-Lens RF lens setting of 60, an AGC target of 1e6, a max injection time of 10ms in profile mode. For dependent scans, an intensity filter of 1e3, charge state selection of 2 – 4 charges, and dynamic exclusion for 15 seconds with 10ppm low and high tolerances were used. A 2m/z window was used prior to CID fragmentation with a setting of 35%. Data acquisition was carried out in the ion trap using the ‘Normal’ scan rate, an AGC target of 1e4, and a max injection time of 100ms in centroid mode.

### Mass spectrometry data analysis

All data files were processed with RawTools as described in the main text. For comparison of RawTools and ProteoWizard^9^ generated MGF output, a combination of SearchCLI (version 3.3.1)^10,11^ and PeptideShakerCLI (version 1.16.23)^12^ was used. All searches used the X!Tandem (version 2015.12.15.2)^13^ algorithm. MS2 data were searched against a UniProt human proteome database (version 2018_09) containing common contaminants (The Global Proteome Machine cRAP sequences - https://www.thegpm.org/crap/) that was appended to reversed sequences generated using the –decoy tag of FastaCLI in SearchCLI (42,190 total sequences, 21,095 target). Identification parameter files were generated using IdentificationParametersCLI in SearchCLI specifying precursor and fragment tolerances of 20ppm and 0.5 Da, carbamidomethyl of cysteine as a fixed modification, and oxidation of methionine and acetylation of protein N-term as variable modifications. Trypsin enzyme rules with a total of 2 missed cleavages allowable was specified.

All SearchCLI results were processed into PSM, peptide, and protein sets using PeptideShakerCLI. Error rates are controlled in PeptideShakerCLI using the target-decoy search strategy to determine false-discovery rates (FDR). Hits from multiple search engines are unified using posterior error probabilities determined from the target-decoy search strategy. Results reports were exported from PeptideShakerCLI using the ReportCLI with numeric values provided to the – reports tag to provide the ‘Certificate of Analysis’, ‘Default Protein Report’, ‘Default Peptide Report’, and ‘Default PSM Report’. All results (PSM, Peptide, Protein) were filtered to provide a final FDR level of <1%. Final mzid files output from PeptideShakerCLI used MzidCLI with the default parameters.

For peptide matching as part of the diagnostic ‘preview’ search functionality of RawTools, the IdentiPy (commit version 0275e13)^14^ search engine was used. A total of 1000 MS2 scans (adjustable via the ‘-N’ flag) were extracted from each raw file and fed into IdentiPy on-the-fly. MS2 data were searched against a UniProt human proteome database (version 2018_09) containing common contaminants (21,095 total target sequences). Decoy proteins were generated on-the-fly by IdentiPy using the ‘reverse’ specification for target-decoy analysis. RawTools automatically reads the instrument configuration and adjusts the mass accuracy settings based on the determined mass analyzer. Left and right precursor accuracy settings were set at 10ppm, with product accuracy at 0.5 Da for the LTQ Velos Orbitrap MS1 and MS2 ion trap data. Carbamidomethylation of cysteine was set as a fixed modification and oxidation of methionine variable. Trypsin enzyme rules with a total of 2 missed cleavages allowable was specified. To enhance speed, the ‘auto-tune’ functionality of IdentiPy is automatically disabled in RawTools. IdentiPy results are filtered by taking the 95^th^ percentile decoy score and keeping all target hits above this value.

### General statistical parameters

In all boxplots, center lines in plotted boxes indicate the median, upper and lower lines the 75^th^ and 25^th^ percentiles, and upper and lower whiskers 1.5X the interquartile range. The calculation of individual p-values was performed using two-sided Students t-tests of sample sets, unless otherwise noted.

### Data and code availability

The mass spectrometry proteomics data were deposited to the ProteomeXchange Consortium (http://proteomecentral.proteomexchange.org) via the PRIDE partner repository^15^ with the dataset identifiers: PXD011070 (fraction subset data), and PXD011069 (diagnostic QC data). The repository contains all raw data, search results, and sequence database files.

R Notebook files detailing data analysis and figure creation for this manuscript are all freely available on the RawTools GitHub page: https://github.com/kevinkovalchik/RawTools/tree/master/documentation/manuscript/Rscripts_for_data-analysis.

## Results and Discussion

RawTools was developed to improve the previously described RawQuant^8^ tool that was used for parsing of scan and quantification data directly from of raw files acquired on Thermo Orbitrap MS instruments. The use of Python programming language made RawQuant installation and use of the tool less user-friendly and ultimately limited the efficiency of raw file processing. In addition, RawQuant was built around the use of the Thermo MSFileReader library, which restricted raw file processing to Windows systems. To improve on these aspects, RawTools has been developed in the ‘.NET’ language C# and was built to utilize the Thermo RawFileReader library. As a result of these changes, RawTools is now distributed as a single, user-friendly package that rapidly processes raw MS data derived from Thermo Orbitrap instruments in an operating system-independent manner (e.g. Windows, MacOS, and Linux).

Like RawQuant, RawTools includes functionality for parsing scan and quantification data from the raw MS files of a variety of instrument architectures (e.g. Q-Exactive and Fusion families – including the HF-X and Lumos, LTQ-Orbitrap family) without any pre-conversion or processing (Figure 1). However, RawTools includes new functionalities to greater facilitate interrogation of parsed scan and quantification data, including: 1. An improved ‘Metrics’ text output that provides a detailed summary information on MS performance, 2. Improved scan ‘Matrix’ text output, including scan-by-scan measures of duty cycle time, number of triggered MS2 scans, precursor ion abundance and elution windows, 3. Automatic linking of identification and quantification scans when using isobaric tag quantification, and 4. Direct output of total or base-peak intensity chromatographic data. As with RawQuant, all processed data are output in simple tab-delimited text files that can be easily input into supplementary analysis tools for visualization (e.g. Excel, Python, R). RawTools can also be used to generate MGF output for use with database matching search tools. Moreover, RawTools now includes additional functionality that enables mass and intensity filtering of the generated MGF output (e.g. to remove isobaric reporter ions prior to data searching). From a quantification standpoint, RawTools incorporates the functionality of RawQuant for the extraction of isobaric tag quantification data directly from raw files (including signal intensity, noise, resolution), while also adding the ability to perform precursor ion abundance analysis (Figure 1).

RawTools also includes a newly developed implementation of a diagnostic ‘QC’ feature that can be used to achieve and maintain optimum instrument performance (Figure 1). The diagnostic QC feature of RawTools uses a combination of the outputs gathered from the parsing of scan and quantification data to compile summary data that can be used to aid in informed decision-making related to instrument performance. In addition, the diagnostic QC analysis includes an integrated ‘preview’ database search using the IdentiPy tool, facilitating calculation and monitoring of a wide collection of identification-related metrics, including: 1. MS2 identification rates, 2. Enzyme cleavage efficiency, and 3. Modification or labeling efficiency. Importantly, due to the exceptional speed of RawTools and the direct processing of the raw output data from an Orbitrap MS, diagnostic QC analysis can be completed in near real-time. As a final feature to aid in the visualization and interpretation of RawTools diagnostic QC results for users of all experience levels, an R Shiny application has been developed that generates high-quality plots of summarized user input data through a publicly accessible web interface.

To demonstrate the functionality of RawTools for data processing and analysis, a single tryptic digest of a HeLa cell lysate was prepared and injected repeatedly (n = 140 individual injections) in 30-minute runs on an Orbitrap Velos MS instrument. The LC-MS system was cleaned and calibrated and a fresh chromatography column prepared and equilibrated prior to the first injection to ensure optimum performance. This design allowed for the direct visualization of the performance degradation of the Velos MS over time. This set of 140 raw files was used to demonstrate the performance and utility of RawTools in a variety of scenarios as described below.

### RawTools enables efficient parsing and analysis of raw Orbitrap MS files

To first demonstrate the basic parsing functionality of RawQuant, the raw files from the first 10 injections of the 140-raw file set were used as a test data set. As these represent the initial injections, these data files should represent the optimum of instrument performance. To benchmark the performance of RawTools, the test set was processed separately on Windows (Core i7-6400 @ 3.4GHz, 32GB of RAM, 64-bit Windows 10) and Linux (Xeon E5-2690 @ 2.9GHz, 132GB of RAM, CentOS 7) systems. Each of the computational setups efficiently processed the entire 10-file data set, requiring just 01m12s and 02m07s (for simplicity, time is given in the notation of hours, minutes, seconds – 00h00m00s) on the Windows and Linux systems, respectively. To put these times in context, on the same Windows machine, RawQuant (version 0.2.3) and ProteoWizard (version 3.0.18225 64-bit) required 2m36s and 1m53s just to generate individual MGF outputs for each of the files across the entire set. However, as part of processing, RawTools was also generating summarized metrics and parsed scan and quantification data files in this same time window along with MGF creation. Compared to its predecessor RawQuant, this represented a 116% speed improvement despite the extraction of additional information by the RawTools software.

During parsing, RawTools can be used to generate two types of text output, Metrics (-x flag in RawTools) and Matrix (-p or -q flag in RawTools, combined with –u for precursor quantification) files. The Metrics files contain summarized information on the MS operation during the acquisition. Using the metrics data, properties such as the scan numbers, rates, duty cycle characteristics, and values relating to chromatography performance (e.g. column peak capacity) can be easily visualized (Figure 2a-f). Interrogation of these values can provide valuable insight into targetable areas for improvement of instrument performance, as demonstrated previously^8^. In the case of the 10-replicate subset examined here, the summarized metrics data illuminated a potential problem with the third replicate. Despite being a repeat of the other runs, the third injection had a reduced MS2 acquisition rate resulting in fewer scans acquired overall (Figure 2). However, although it was clear something was wrong with this replicate, it was not immediately obvious from the summarized metrics exactly what the issue was.

**Figure 2.**
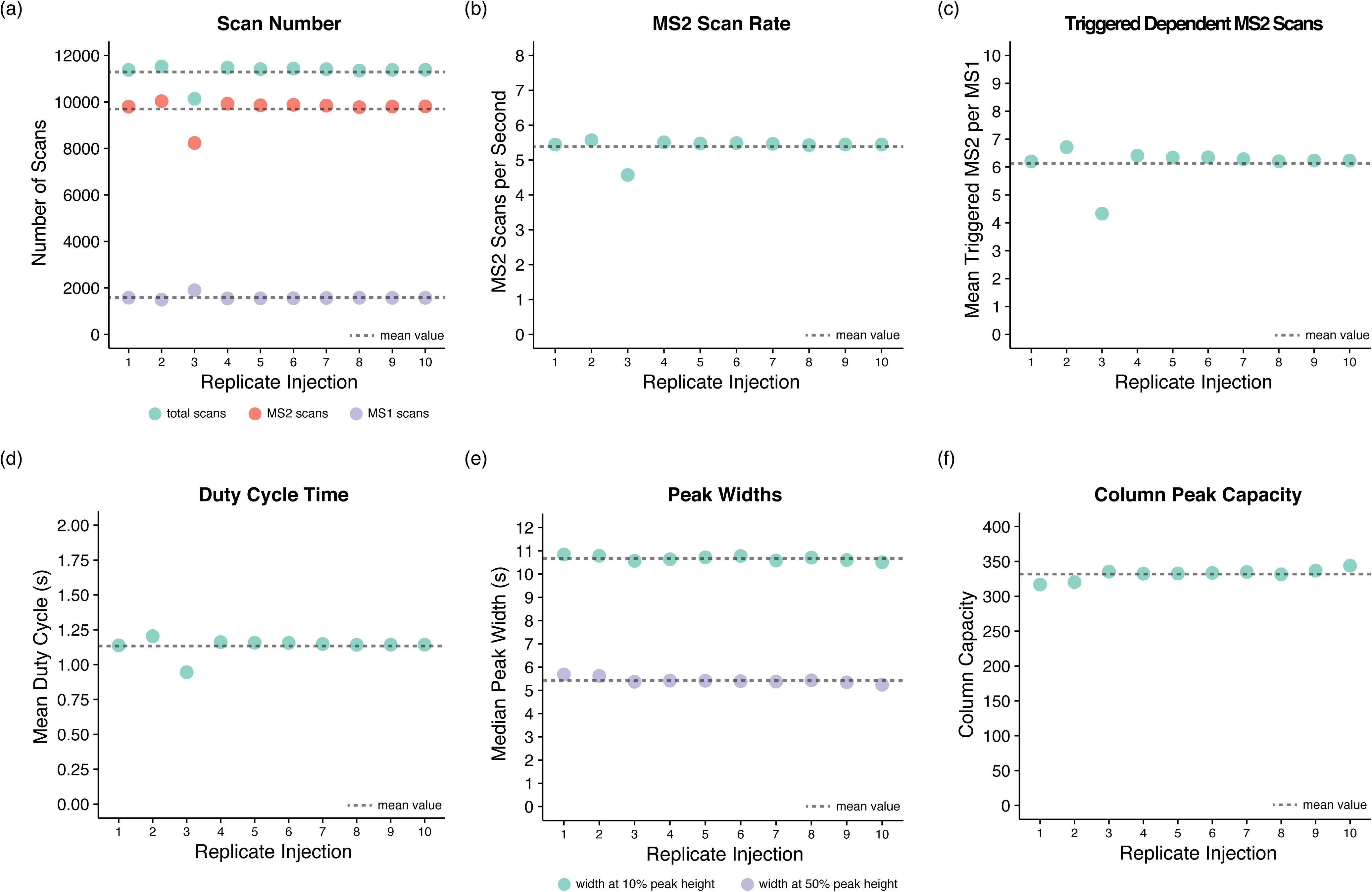
RawTools enables rapid and dynamic analysis of raw data files to illuminate MS performance. A subset (n = 10) of raw files acquired as part of a replicate injection set derived from a HeLa tryptic digest were analyzed with the RawTools parse functionality to generate ‘Metrics’ files. The resultant text output from RawTools was investigated to generate insights into: **(a)** Scan numbers, **(b)** MS2 scan rates, **(c)** Numbers of dependent scans triggered per MS1, **(d)** Duty cycle duration, **(e)** Chromatographic peak width, and **(f)** Column peak capacity. Dashed lines on each plot indicates the mean across the 10 replicate injections for the displayed values.

To investigate the third injection further, the Matrix output of RawTools was used. When RawTools is invoked with the ‘-p’ flag, the Matrix output contains individual information for all scan events contained within the raw MS file. Alternatively, when used with the ‘-q’ flag, the scan Matrix output from RawTools additionally contains information related to isobaric tag quantification. With the Matrix output from replicate 3 of the subset, plotting of the MS2 acquisition rate and MS1 parent ion intensities across the entire acquisition window revealed an unexplained gap in the electrospray in the early stages of the gradient, lasting just 5 minutes (Figure 3a-b). This short drop in spray resulted in no triggered of MS2 scans, but did not impact the overall quality of chromatography as measured by peak shape and column capacity (Figure 2e-f). These data highlight the utility of measuring multiple metrics of instrument performance, and the ability of RawTools to facilitate easy illumination of potentially sub-optimal MS runs.

**Figure 3.**
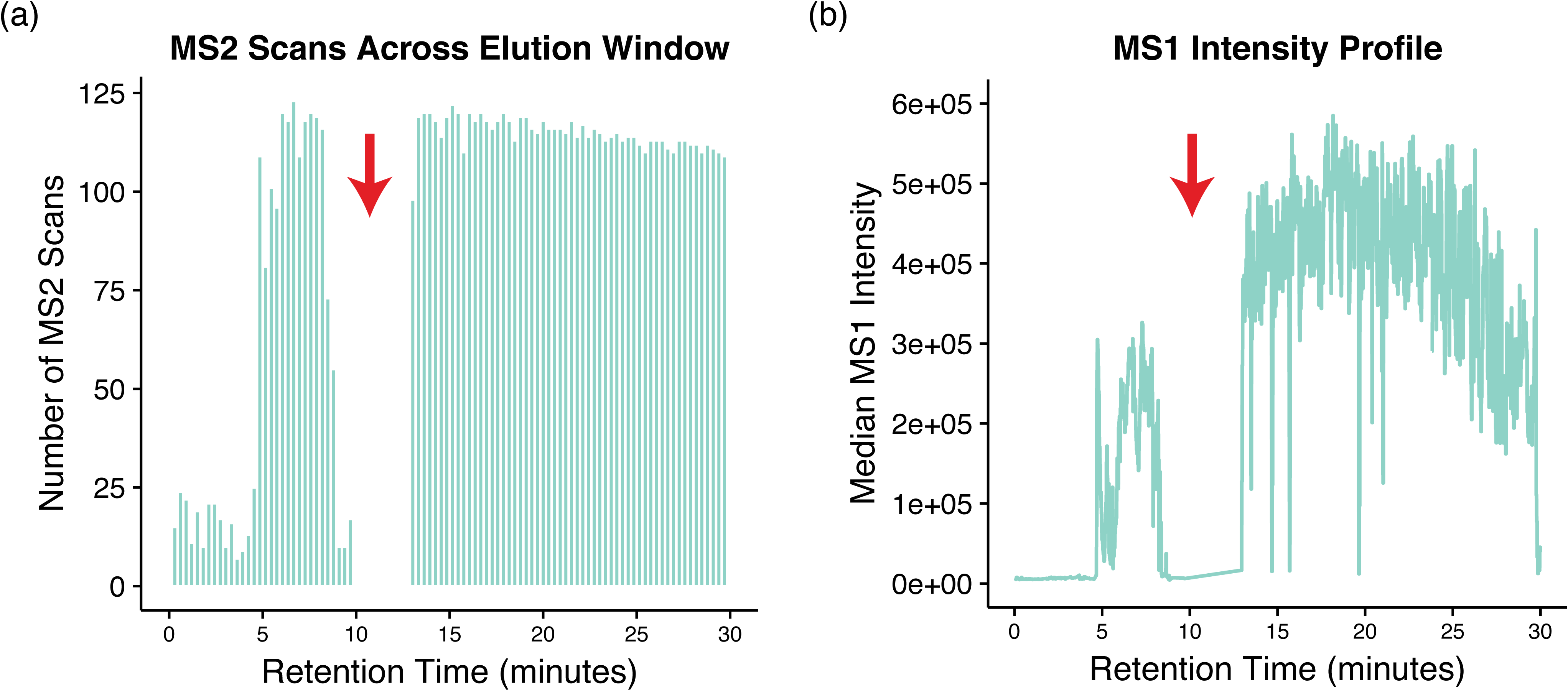
RawTools enables simplified detection of errors that occur during MS acquisition. The third injection from the 1 – 10 injection set was examined further using the RawTools parse functionality to generate scan ‘Matrix’ files to determine the cause of the difference in relation to the other replicate samples. The resultant text output from RawTools was used to identify a break in the spray being generated from the nanospray tip, as observed as a gap in **(a)** MS2 scan acquisition and **(b)** Intensity of precursors in MS1 scans.

Lastly, to establish the validity of the MGF output from RawTools, the results from a database search was carried out and the results compared with a file generated using the established ProteoWizard software tool. The MGF outputs for the 10-replicate subset from RawTools and ProteoWizard were individually searched using SearchCLI and PeptideShaker. Comparison of the unique peptide and protein identification rates (1% FDR) revealed no significant differences between the results when using MGF outputs from the individual software tools (p = 0.99 for identification numbers of peptides and proteins, 99% overlap in identifications for both peptides and proteins between RawTools and ProteoWizard sets) (Figure S1). Taken together, these data demonstrate the effective parsing and extraction of meta and scan data from Thermo Orbitrap raw files using RawTools towards general identification analyses or in-depth examination of MS performance.

### RawTools facilitates robust tracking of acquisition performance

RawTools includes a newly developed diagnostic ‘QC’ feature designed for the visualization of instrument performance. In addition to providing summarized performance metrics as before (e.g. scan numbers and rates, chromatography performance), RawTools diagnostic QC output currently includes other supplementary data: 1. Electrospray stability, 2. Gradient elution performance, and 3. MS2 fragment signal distributions. In addition, RawTools can employ the IdentiPy search tool to provide a ‘preview’ functionality that enables tracking of data in QC that includes: 1. MS2 identification rate, 2. Modification efficiency (e.g. labeling efficiency), and 3. Analyzer mass error. The output of RawTools QC is a single comma separated file containing compiled data from all files processed to date. Newly acquired files can be appended to this compiled file by simply placing them in a target directory and re-triggering the RawTools QC command. Although this operation is not true real-time because it is not automatically scanning over a directory to monitor for newly generated data, this near real-time implementation allows users greater control over which files are included in a diagnostic QC set and the ability to carry out these tasks in a remote location. Visualization and interpretation of these data are further facilitated via the freely available RawTools R Shiny application web interface.

To demonstrate the performance and utility of this QC feature, the entire 140-raw file set was processed with RawTools on a Linux system (Xeon E5-2690 @ 2.9GHz, 132GB of RAM, CentOS 7). Processing of the entire 140-raw file set for diagnostic QC information required 24m52s without the database search, and 3h36m57s with IdentiPy analysis (01m30s per file, auto-tune disabled, n = 1 variable modification, n = 1000 MS2 spectra) on the Linux system. Although this processing time appears to be substantial, the majority of this time is devoted to the IdentiPy search. For comparison, performing IdentiPy analysis independent of RawTools requires 03h18m35s for the entire 140 raw file set (01m25s per file, auto-tune disabled, n = 1 variable modification, n = 1000 MS2 spectra). Therefore, the RawTools component of the analysis required just 18-minutes (8% of the total processing time). The processing speed when using the ‘preview’ search functionality could be improved in future iterations of RawTools using rapid search engines like X!Tandem^13^.

Investigation of the scan totals calculated by RawTools across the 140-injection set illuminated a gradual but steady decrease in the numbers of acquired MS2 events as the injections continued (Figure 4a). Using the values for the summed signal intensities from MS1 events within each raw file, a continual decrease was again observable, but with specific outliers appearing (Figure 4b). Using the injection 23 outlier as an example for further investigation, the RawTools chromatogram output of this file revealed inconsistencies in the base peak intensity patterns, indicative of unstable electrospray (Figure 4c). These spray instability events were also easily observed based on the electrospray stability output of RawTools (Stability = Number of MS1 scans whose neighbor differs in signal by >10-fold) (Figure S2). Although minor in terms of duration and frequency, these events made enough of an impact to be flagged as potentially problematic via user interpretation of the generated RawTools output.

**Figure 4.**
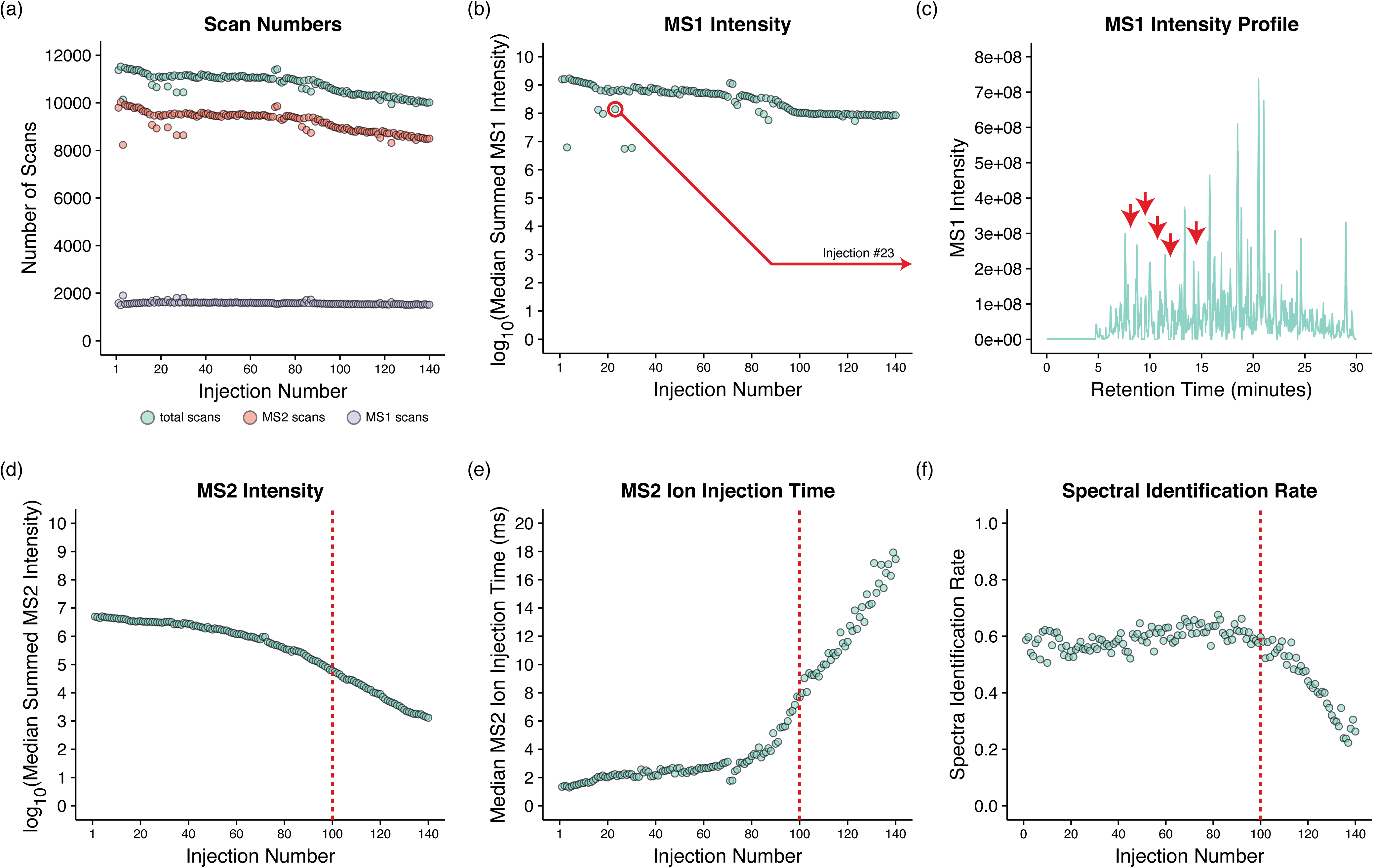
RawTools QC analysis facilitates illumination of variation in MS operational performance. The entire set of (n = 140) HeLa replicate injections was analyzed with the diagnostic QC feature of RawTools to generate a single comma separated summary output. The resultant data were used to probe **(a)** Scan numbers and **(b)** MS1 intensities to reveal inconsistencies. **(c)** Selected base peak chromatogram of MS1 intensities demonstrating spray instability. Red arrows in the mark areas of electrospray instability. The RawTools data were further examined to highlight instrument performance degradation via **(c)** MS2 intensities, **(d)** MS2 injection times, and **(e)** IdentiPy MS2 spectral identification rates. Dashed red lines indicate the 100^th^ sample injection.

As expected, across the 140 injections the steady decline in MS1 signal was indicative of a loss of sensitivity in the MS, a trend that is easily visible in both the decrease in summed signals observed in MS2 scans, as well as the increasing ion injection times required to hit the same specified ion target value (Figure 4d-e). This drop in MS2 quality is reflected in the observed decrease in peptide identification rates as determined with the ‘preview’ search done with IdentiPy (Figure 4f). This drop in spectral identification rate may also include a contribution by the observed drift in the mass accuracy of the mass analyzer used for MS1 precursor acquisition across the replicate injections (Figure S3). RawTools also enabled identification of areas where stability is observed, such as sample preparation metrics like enzyme digestion efficiency and the proportions of oxidation events on methionine-containing peptides (Figure S4a-b). Although not demonstrated here, the modification frequency calculations performed by RawTools can be used to track properties such as labeling efficiency when using isobaric tags. Altogether, these results demonstrate the utility of RawTools for processing data in a QC setting towards longitudinal tracking of instrument and experiment performance, and highlights the ability to identify even minute disruptions in expected MS operation.

In this work, the development and use of the RawTools software tool is described. RawTools is a simplified and streamlined version of the previously presented RawQuant tool that now provides improved flexibility in implementation, additional data outputs, and a QC feature for performance tracking. Importantly, RawTools operates directly on the raw output of Thermo Orbitrap MS instruments with no additional data conversion. Substantial effort has gone into simplifying RawTools and the output from the tool in order to improve usability for users of all skill levels. As an example of this, the RawTools R Shiny application provides users with a simple point-and-click interface through which their QC data can be visualized. It is also noteworthy that the majority of the plots used in this work were generated using code from the RawTools R Shiny application. Although not discussed in detail here, RawTools has substantial utility in the optimization of MS method acquisition parameters and in the extraction of quantification results, directly from raw MS data as established previously^8^. Importantly, all the data discussed here are available almost immediately after MS analysis due to the exceptionally rapid processing times of RawTools, the ability to work directly with raw Orbitrap files, and the ease with which the output data can be handled via the provided web-application. Taken together, the improved design and newly implemented features of RawTools position it as a powerful software for interrogation of MS operation dynamics and performance.

## Supporting Information

The following files are available free of charge at ACS website http://pubs.acs.org:

- Supporting information describing the equivalency in MGF output between software tools (Figure S1). Information relating to the measured instability of electrospray during extended MS analysis (Figure S2) and the increase in mass detection error (Figure S3). Information relating to the monitoring of the stability in sample preparation using digestion efficiency and methionine oxidation frequency (Figure S4).

## Acknowledgements

C.S.H. would like to acknowledge valuable input and discussions from Lida Radan and Philipp Lange. All authors would like to acknowledge valuable support from the Genome Sciences Centre Systems department, specifically Ross Stevenson and Brendan O’Huiginn for their help with hosting the R Shiny web application.

## Funding Sources

This work was supported by the British Columbia Cancer Foundation (G.B.M., C.S.H, S.M.) and a Discovery Grant from the Natural Sciences and Engineering Research Council (NSERC) of Canada (G.B.M. and D.D.Y.C.). K.A.K. acknowledges Four-Year Doctoral Fellowships from the University of British Columbia (award numbers - 6569, 6456) and a Gladys Estella Laird Research Fellowship (award number 4818).

## Competing Financial Interests

The authors declare no competing financial interests.

## Author Contributions

K.A.K. and C.S.H. conceived the idea, carried out the data analysis, and wrote the manuscript. S.C. and S.S. helped with data acquisition and tool design. P.H.S., D.D.Y.C., and G.B.M. contributed to writing of the manuscript.

## References

(1) Nesvizhskii, A. I. A Survey of Computational Methods and Error Rate Estimation Procedures for Peptide and Protein Identification in Shotgun Proteomics. J. Proteomics 2010, 73 (11), 2092–2123.

(2) Bantscheff, M.; Lemeer, S.; Savitski, M. M.; Kuster, B. Quantitative Mass Spectrometry in Proteomics: Critical Review Update from 2007 to the Present. Anal. Bioanal. Chem. 2012, 404 (4), 939–965.

(3) Bittremieux, W.; Valkenborg, D.; Martens, L.; Laukens, K. Computational Quality Control Tools for Mass Spectrometry Proteomics. Proteomics 2017, 17 (3–4).

(4) Bittremieux, W.; Tabb, D. L.; Impens, F.; Staes, A.; Timmerman, E.; Martens, L.; Laukens, K. Quality Control in Mass Spectrometry-Based Proteomics. Mass Spectrom. Rev. 2018, 37 (5), 697–711.

(5) Rudnick, P. A.; Clauser, K. R.; Kilpatrick, L. E.; Tchekhovskoi, D. V.; Neta, P.; Blonder, N.; Billheimer, D. D.; Blackman, R. K.; Bunk, D. M.; Cardasis, H. L.; et al. Performance Metrics for Liquid Chromatography-Tandem Mass Spectrometry Systems in Proteomics Analyses. Mol. Cell. Proteomics MCP 2010, 9 (2), 225–241.

(6) Amidan, B. G.; Orton, D. J.; Lamarche, B. L.; Monroe, M. E.; Moore, R. J.; Venzin, A. M.; Smith, R. D.; Sego, L. H.; Tardiff, M. F.; Payne, S. H. Signatures for Mass Spectrometry Data Quality. J. Proteome Res. 2014, 13 (4), 2215–2222.

(7) Trachsel, C.; Panse, C.; Kockmann, T.; Wolski, W. E.; Grossmann, J.; Schlapbach, R. RawDiag: An R Package Supporting Rational LC-MS Method Optimization for Bottom-up Proteomics. J. Proteome Res. 2018, 17 (8), 2908–2914.

(8) Kovalchik, K. A.; Moggridge, S.; Chen, D. D. Y.; Morin, G. B.; Hughes, C. S. Parsing and Quantification of Raw Orbitrap Mass Spectrometer Data Using RawQuant. J. Proteome Res. 2018, 17 (6), 2237–2247.

(9) Chambers, M. C.; Maclean, B.; Burke, R.; Amodei, D.; Ruderman, D. L.; Neumann, S.; Gatto, L.; Fischer, B.; Pratt, B.; Egertson, J.; et al. A Cross-Platform Toolkit for Mass Spectrometry and Proteomics. Nat. Biotechnol. 2012, 30 (10), 918–920.

(10) Vaudel, M.; Barsnes, H.; Berven, F. S.; Sickmann, A.; Martens, L. SearchGUI: An Open-Source Graphical User Interface for Simultaneous OMSSA and X!Tandem Searches. Proteomics 2011, 11 (5), 996–999.

(11) Barsnes, H.; Vaudel, M. SearchGUI: A Highly Adaptable Common Interface for Proteomics Search and de Novo Engines. J. Proteome Res. 2018, 17 (7), 2552–2555.

(12) Vaudel, M.; Burkhart, J. M.; Zahedi, R. P.; Oveland, E.; Berven, F. S.; Sickmann, A.; Martens, L.; Barsnes, H. PeptideShaker Enables Reanalysis of MS-Derived Proteomics Data Sets. Nat. Biotechnol. 2015, 33 (1), 22–24.

(13) Craig, R.; Beavis, R. C. TANDEM: Matching Proteins with Tandem Mass Spectra. Bioinforma. Oxf. Engl. 2004, 20 (9), 1466–1467.

(14) Levitsky, L. I.; Ivanov, M. V.; Lobas, A. A.; Bubis, J. A.; Tarasova, I. A.; Solovyeva, E. M.; Pridatchenko, M. L.; Gorshkov, M. V. IdentiPy: An Extensible Search Engine for Protein Identification in Shotgun Proteomics. J. Proteome Res. 2018, 17 (7), 2249–2255.

(15) Vizcaíno, J. A.; Csordas, A.; Del-Toro, N.; Dianes, J. A.; Griss, J.; Lavidas, I.; Mayer, G.; Perez-Riverol, Y.; Reisinger, F.; Ternent, T.; et al. 2016 Update of the PRIDE Database and Its Related Tools. Nucleic Acids Res. 2016, 44 (22), 11033.

